# Feasibility of clinician-facilitated 3D printing of synthetic cranioplasty flaps

**DOI:** 10.1101/251488

**Authors:** Sandip S Panesar, Joao Tiago A Belo, Rhett N D’Souza

**Author notes:** Corresponding Author: Sandip S Panesar, Department of Neurological Surgery, 200 Lothrop Street, Pittsburgh 15213, Pennsylvania, United States of America. **Funding:** No applicable funding.

## Abstract

**Objectives:** 3D scanning and stereolithographic printing technology becoming increasingly common, however its implementation into clinical practice is in its primacy. These technologies may be esoteric to the practicing neurosurgeon. We explored a range of 3D scanning and stereolithographic techniques to create patient-specific synthetic implants.

**Methods:** We simulated bilateral craniectomies from a single cadaveric specimen to create 3 methods of creating stereolithographically-viable virtual models. Firstly, we used ‘pre-and-post operative’ CT derived bony windows to create a virtual skull model, from which the flap was extracted. Secondly, we used an entry-level 3D light-scanner to scan and render models of the individual bone pieces. Thirdly, we used an arm-mounted, 3D laser-scanner to create virtual models using a real-time approach.

**Results:** Flaps were printed from the CT scanner and laser scanner models only, in a UV-cured polymer. The light scanner did not produce suitable virtual models for printing. The CT scanner derived models required extensive post-fabrication modification to fit the existing defects. The laser-scanner models assumed good fit within the defects without any modification.

**Conclusions:** The methods presented varying levels of complexity in acquisition and model rendering. Each technique required hardware at varying in price points from $0 to ∼$100,000. The laser-scanner models produced the best quality parts which bore near-perfect fit with the original defects. We discuss potential neurosurgical applications of this technology.

## 1. Introduction

Cranioplasties are indicated for filling bony skull defects to serve cosmetic or medical reasons (Andrabi, Sarmast, Kirmani, & Bhat, 2017; Dujovny et al., 1997; Goldstein, Paliga, & Bartlett, 2013). Skull defects may be traumatic or post-surgical, and can vary in size and shape. Often the region requiring replacement is of compound, 3-dimensional (3D) shapes, and involves multiple constituent bones of the cranial vault. In certain cases, replacement with the removed bone may be unfeasible, such as in traumatic head injuries with compound skull fractures and for emergency craniectomies, where the time between craniectomy and cranioplasty may be considerable (Flannery & McConnell, 2001). Aside from a physical barrier to trauma, replacement of a bony skull defect can aid cerebrospinal fluid dynamics (Beauchamp et al., 2010; Dujovny et al., 1997), act as a barrier to pathogens (Beauchamp et al., 2010) and serves a cosmetic purpose (Dujovny et al., 1997; Rotaru et al., 2012). A variety of graft procedures and materials have been developed to tackle cranioplasty requirements (Aydin, Kucukyuruk, Abuzayed, Aydin, & Sanus, 2011; Zanotti et al., 2016). The material may be harvested from the patient (bone, fat or tissue), cadavers or animals. Synthetic cranioplasties were first utilized following the introduction of mass-produced synthetic polymers and pioneering reconstructive surgery necessitated by the World Wars (Harris et al., 2014). Modern synthetic materials for alloplastic cranioplasty include methacrylate, polyether ether ketone (PEEK), silicon, ceramics and titanium (Aydin et al., 2011). When choosing synthetic allografts surgeons must consider not only the physical properties of the material including shaping, strength and fixation (Ridwan-Pramana et al., 2017), but also its biostability (Gautschi, Schlett, Fournier, & Cadosch, 2010; Kim et al., 2013). Cranioplasty implant methods are therefore dependent on the choice of material, size of bony defect and fixation. They may include taking molds of the defect or from the removed bone (Fathi, Marbacher, & Lukes, 2008) or using a foundational titanium mesh to support overlying synthetic material (Malis, 1989; Ng, Ang, & Nawaz, 2014), amongst others.

3D printing or stereolithography is now a widely accepted method of creating complex physical objects out of plastic or metallic materials using a virtual template and a stereolithographic printer. For templates, reconstruction of the bony component of computerized tomography (CT) scans may be utilized to demonstrate the dimensions and shape of the defect (Barker, Earwaker, & Lisle, 1994; Bouyssie, Bouyssie, Sharrock, & Duran, 1997). In-house software routinely used for clinical image viewing such as Picture Archiving and Communication System (PACS) (Arenson, 1992) allows clinical viewing of the radiographic images, and potentially 3D reconstructions of various tissue-windows. In order to manipulate the data, however, it is likely that third-party software will be required. This software should permit reconstruction of the standardized Digital Imaging and Communication in Medicine (DICOM) (Ratib, Ligier, & Scherrer, 1994) data within a 3D space. It should be able to manipulate the models and export it as a stereolithography (.stl) or object (.obj) file for use by 3D computer-aided design (CAD) solutions. Various commercial and industrial 3D manufacturing devices and techniques are available at varying price points and technical requirements (Gebhardt, 2012). Thus far, the technology has primarily been used for medical modelling and training purposes (McGurk, Amis, Potamianos, & Goodger, 1997; Rengier et al., 2010). Despite growing awareness amongst clinicians, the technology has not been widely implemented in neurosurgery (Baskaran, Strkalj, Strkalj, & Di Ieva, 2016; Jimenez Ormabera et al., 2017; Klein, Lu, & Wang, 2013; Randazzo, Pisapia, Singh, & Thawani, 2016; Tomasello, Conti, & La Torre, 2016; Weinstock et al., 2017). This may be due to lack of awareness, cost of equipment, perceived learning curves and importantly, the relative lack of FDA-approved printing mediums suitable for human implantation.

The purpose of this paper is therefore to demonstrate simplified methods of creating synthetic cranioplasty flaps using minimal specialized software and with commercially available hardware. The rationale for our study is to integrate non-clinical advances in 3d-image acquisition, clinical software and 3D printing technology with the hope that our methods may be employed in the future to create clinically viable synthetic cranioplasty allografts by clinicians on-site and with minimal prior training.

## 2. Methods

### 2.1 Cadaveric Specimen

2 craniectomies were simulated on a cadaveric specimen. A right-sided temporal craniectomy (47 mm maximum anteroposterior length x 51 mm maximum vertical height) to mimic a middle fossa approach, and a left-sided hemicraniectomy (141 mm maximum anteroposterior length x 108 mm maximal vertical height) to mimic a decompressive procedure. Once the craniectomies had been performed, bone flaps were carefully separated from adherent dura and freed from the skull. Dura and periosteum were then removed manually. There were several metallic nails placed within the specimen, intended for fiducial navigation, these were removed from the pieces and holes filled with a 2-part epoxy resin. Along its superior orbital, frontal, parietal and occipital extremities, it had a thickness of ∼ 3 mm, narrowing to <1 mm thickness at its inferior temporal portion. The smaller piece was flat and rhomboid-shaped, with a uniform thickness of ∼ 3 mm. Along its edges, it had a thickness of ∼ 3 mm, narrowing to <1 mm thickness at its inferior temporal portion.

### 2.2 Computerized Tomography Method

The specimen received both pre-and-post-craniectomy fine-cut CT scans on a GE VCT 64 Slice scanner (GE Healthcare, Chicago, U.S.A.). Scan type was helical, with a rotation time of 0.6 seconds, a detector coverage of 40mm, a pitch of 0.516:1, speed 20.62, slice thickness 1.25mm x 1.25mm. Scan field of view was at the head setting, the matrix size was 512, recon type was standard with a full recon option. Tube voltage was 120 and mAs was 280. Window length/window view was 400/40. After acquisition, raw DICOM data from each scan was imported into free 3D Slicer software (www.slicer.org). The bony windows of the scans were rendered as 3D objects and exported as .stl files. Initial editing of the data was conducted using DAVID 3D software (HP Inc., Palo Alto, U.S.A.). Larger tomographic artifacts were manually removed from both scans. After cleaning, the pre-and-post-operative reconstructions were assigned different colors and automatically aligned with each other. To achieve an outline of the craniectomy defect, the post-operative reconstruction was offset within the lateral plane by 0.5mm. This created a suitable impression of the craniectomy defect superimposed upon the pre-operative reconstruction, and the margins of the craniectomy were manually traced on the intact model (Figure 1A). The surroundings of both pre-and-post-operative renders were deleted in their entirety, leaving only the traced flap portion. As this object consisted of an interior surface, and an exterior surface, the fill tool within the software was used to digitally close the defects created by the removal of the overlying skull (Figure 1B-C). The completed render was then saved as a .stl file prior to printing. Total time for creating both models, following scans, was ∼ 2 hours.

**Figure 1 A-C.**
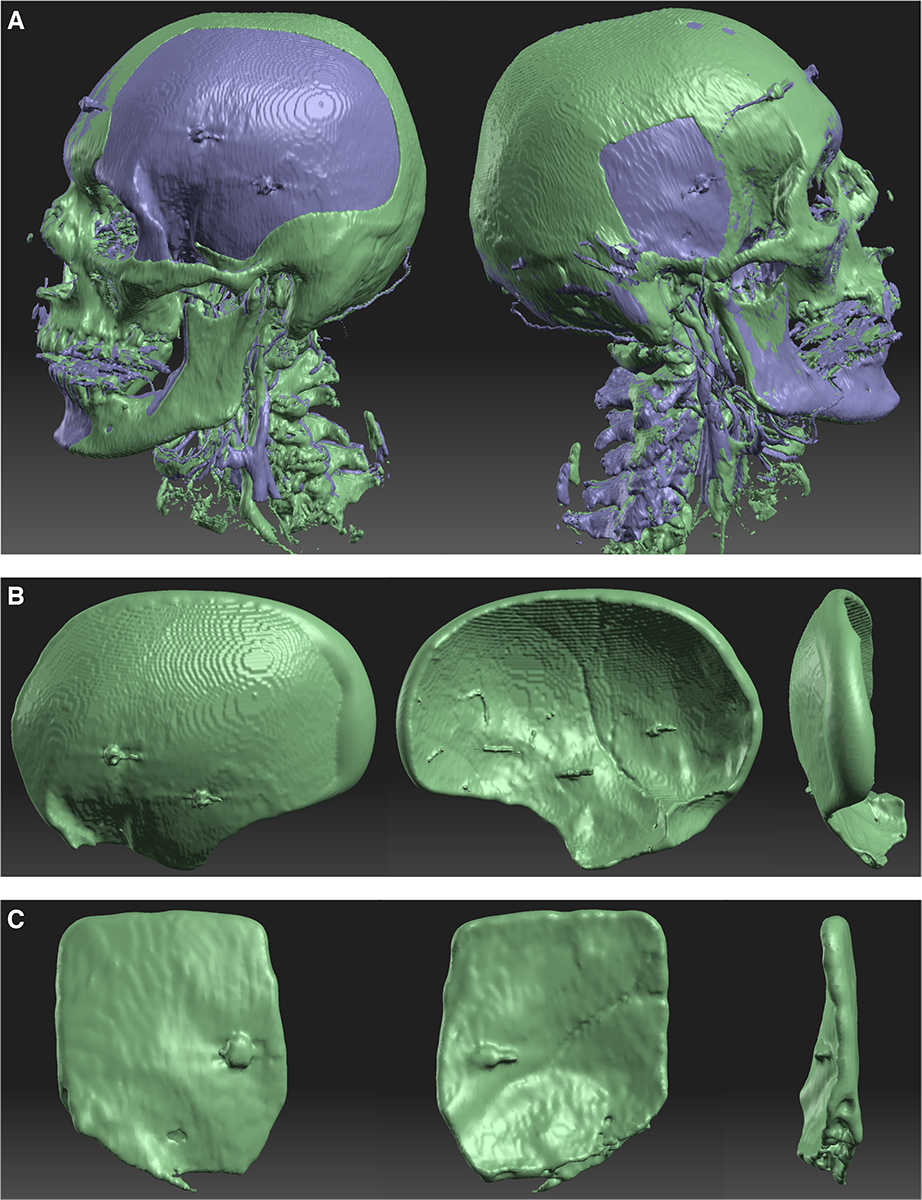
A – Right and left sided views of both CT-derived 3D skull models aligned, and offset to create an impression of the craniectomy defect, and prior to manual tracing and selection. B – Left-Right: Outer, inner and posterior views of the hemicraniectomy model derived from CT scan. Discernible are the concentric layers representative of the 1.25mm CT slicing. There is a prominent raised lip on the exterior, posterior surface, which continues around the edges of the piece. Thickness is best appreciated from the far-right view. These edges are discernible on the middle image and are a result of automated closure of the gap created between interior and exterior surfaces during model construction. The interior surface contains a well-defined middle-meningeal groove. Other features include a portion of the orbito-frontal bone and 2 metallic-screw artefacts. The inner surface contains minor vascular artefacts created as a result of the cadaveric fixation process, in addition the aforementioned screws. C – Left-Right: Outer, inner and anterior views of the middle-fossa approach flap. The piece is approximately rhomboidal in shape. CT-derived tessellation artefacts are not as prominent compared to figure B. Again, a metallic screw artefact is prominent on both exterior and inner surfaces. A portion of reproduced bony pneumatisation is visible on the ventral aspect of the far-right image.

### 2.3 Tripod-mounted Light Scanner

We used the DAVID SLS 3 system, with 4M model camera (HP Inc., Palo Alto, U.S.A.), mounted on a tripod. Briefly, the camera specifications include an Acer K132 (1280 x 800 @ 60 frames per second) projector (Acer Inc., Hsinchu, Taiwan) with a frame limit of 60 frames per second. The bone pieces were separately scanned by placing them upon the supplied turntable, with a 360° range of motion. Prior to scanning, the camera was calibrated according to the size of the bones, using supplied calibration apparatus. The larger hemicraniectomy was calibrated at the 120mm scale, and the smaller piece at the 60mm scale. The camera exposure and projector brightness were automatically controlled and varied depending upon the object position. During the scanning process, 66 patterns were projected for each acquisition. Smoothing average was set to 0, quality check setting was set at 0.5 and outlier removal at 0.1%. The hemicraniectomy required 18 individual scans and the smaller piece 13. Each component scan was then analyzed and automatically aligned to create a complete 3D model. The completed models were directly saved as .stl files. Total scanning time and model editing time for the hemicraniectomy was ~ 4 hours, and ~ 3 hours for the smaller piece (Figure 2A-B).

**Figure 2 A-B.**
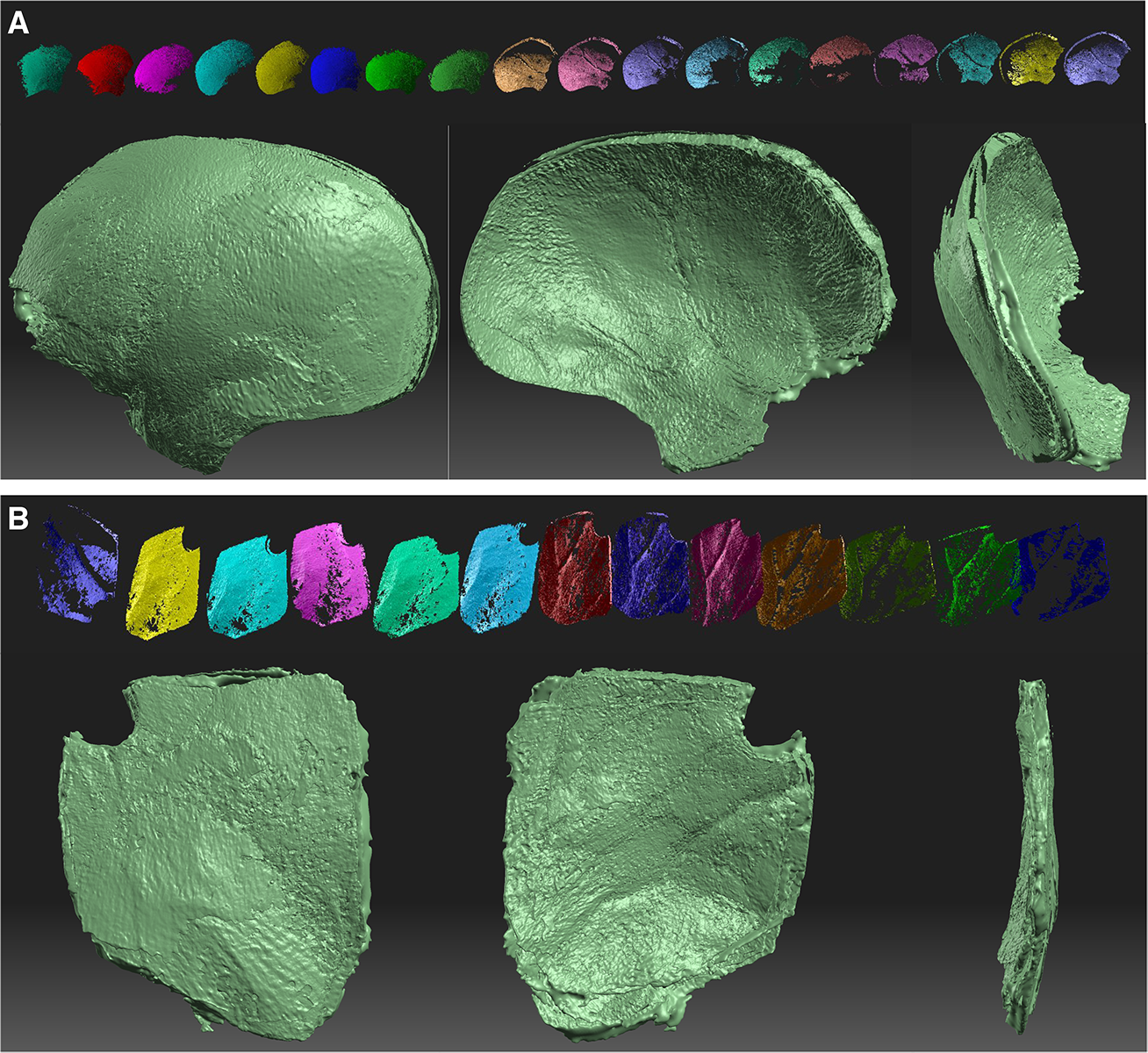
A – Left-right: Outer, inner and posterior views of the light-scanner derived hemicraniectomy model. On top, from left-right are the 18 individual composite images that comprised the final model. Notable on all images is the lack of vascular or metallic artefacts, as the screws were removed prior to scanning. All 3 images demonstrate a non-uniform B – Left-right: Outer, inner and anterior views of the light scanner derived middle-fossa approach model. On top, from left-right are the 13 individual composite scans that were used to compile the final model. Like the larger piece, surface representation is sub-optimal and not representative of the real specimen. Furthermore, scans are poorly aligned, despite automated alignment features, as visible around the edges of all pieces.

### 2.4 Arm-mounted Laser Scanner

We used the FARO Laser ScanArm (FARO Technologies, Lake Mary, U.S.A.) for laser scanning of the removed skull pieces. This is a hand-held ‘gun’ shaped device attached to an articulated arm, allowing free-floating movement in 7 axes. It allows scanning of objects either by contact or at a distance using a mounted laser. We used the latter. It is accurate to a distance of 50μm, with a repeatability of 50μm, 2σ. Its standoff distance is 95mm and has a depth of field of 85mm. Scan width is 34mm at close distances and 60mm at the range. We used its default scan rate of 30 fps x 640 points per line, giving a total acquisition capacity of 19200 points per second. GeoMagic Studio 2012 (GeoMagic, Morrisville, U.S.A.) was used to acquire and compile .stl files in real time. Each piece of bone was mounted upright in a clamp, whilst a continuous scan of inner and outer surfaces was acquired as a single capture, and with the camera held ∼ 9cm from the bone surface. Each bone was rotated once in the clamp to acquire the portion that was obscured by the apparatus during the initial scan. A total of 2 component scans comprised each model. Virtual model creation from the two scans was continuous and real-time, requiring no manual manipulation or reconstruction to create the final .stl model. Total scanning (and model creation) time for each piece of bone was less than 2 minutes (Figure 3A-B).

**Figure 3 A-B.**
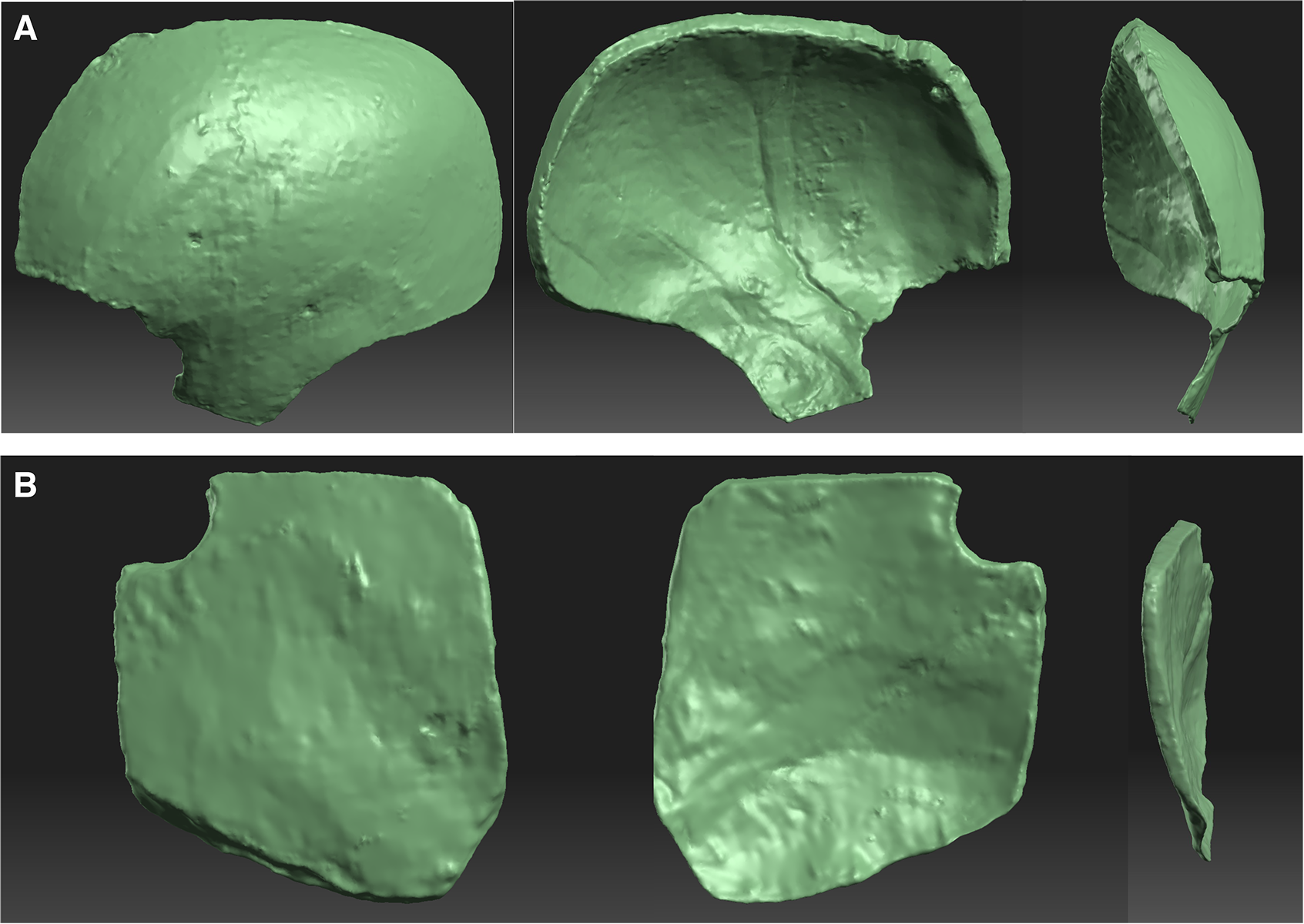
A – Left-right: Outer, inner and anterior views of the laser-scanner derived model. These models provided optimal recreation of the original pieces in terms of size, edges, shape and surface. Note holes where fiducial screws were removed and filled. Furthermore, the pterion is visible on the far-left piece, as are the coronal and squamous sutures. The inner piece contains not only the middle meningeal groove, but also various smaller vascular impressions. On the far-right image, the crispness of edges is clearly visible. B – Left-right: Outer, inner and anterior views of the smaller middle-fossa approach. On the outer aspect, a small indentation left by the removal and filling process of the fiducial screw is visible. On the inner aspect, middle meningeal groove is also visible, however less well pronounced than on the opposite piece, nevertheless, it is discernible on the anterior, head on view of the far-right view.

#### 2.4.1 3D Printing of Virtual Models

As this is a research study into the feasibility of rapidly constructed 3D printed neurosurgical implants we elected to use a plastic medium to mimic commonly used, FDA-approved synthetic polymers. Other considerations were 3D printing equipment available to us, and cost. The flaps were printed using the Viper SLA system (3D Systems, Valencia, U.S.A.). It uses a solid-state laser to facilitate curing of polymer resins during the stereolithographic process. For printing medium we used DSM Somos WaterShed XC 11122 photopolymer (DSM Functional Materials, Elgin, U.S.A.). It is a low-viscosity photopolymer, which allows 3D printing of hard, translucent plastic parts following ultraviolet curing. It has previously been used for medical modeling. Upon further analysis of the light-scanner derived 3D models, we became aware of numerous alignment defects in both the hemicraniectomy and middle-fossa approach models (see 3.1), which rendered them unfeasible for stereolithographic printing without significant and time-consuming editing. We opted to print only the CT-derived and laser-scanner derived models for further comparison. The hemicraniectomy/middle-fossa flaps were printed together during the same process. Total printing time was 24 hours including UV hardening and cleaning for the CT and laser models, respectively.

## 3. Results

### 3.1 Virtual Models

Prior to 3D printing, we compared the quality of the virtual models that were created using each of the 3 construction methods and compared them to the removed cadaveric bone flaps. We used three criteria: Size, shape, edges, thickness and surface quality. In terms of dimensions, all three models all approximated the size of the original pieces, we assessed aligning them within a single space. There were notable variations in edges between the three, particularly between each 3D scanned models and the CT reconstruction. The CT reconstruction of the large hemicraniectomy featured a prominent ‘lip’ around its edges on its inner and outer aspects. The lip was pronounced at the external temporo-occipital surface and occurred as a result of computer-aided filling of the oblique defect between exterior and interior aspects. Elsewhere, the lip was less pronounced as the bone was comparatively thinner. The CT reconstruction of the smaller craniectomy flap was more congruent with the specimen and the 3D scanned models. In terms of thickness, the laser-derived 3D models of the hemicraniectomy and smaller flap were again the most similar to the real bone. Despite using optimized alignment of the composite scans, light-scanner derived models had numerous small defects throughout the entire model. We used the groove of the middle meningeal artery, present on both right-and-left sided flaps to assess surface quality, amongst other textural landmarks. The laser-scanner 3D model appeared to retain the greatest level of surface anatomical detail and overall resemblance, followed by the CT scan-derived model, despite its prominent posterior-external lip. The surface of the light-scanner model appeared grainy and non-uniform. This surface texture was not present on the original specimen (Figures 1–3).

### 3.2 Printed Models

We again used the criteria of size, shape, edges, thickness and surface detail to assess the overall quality of the printed parts. The 3D scanner derived models were identical in dimensions and shape to the specimen flaps. The hemicraniectomy was 139mm (2mm smaller (-1.4%) than specimen) in anteroposterior length and 107 mm (1mm smaller (-0.92%) than specimen) in vertical height. At its thickest (anterosuperior) point, it measured 7mm in thickness, and at its most ventral (temporal section), it was <1mm in thickness. Notably, in this region, there was a small ∼5mm defect within the plastic due to the real specimen being thinner than the minimal thickness reproducible by the printer. Its edges assumed almost identical approximation of the original specimen, with ridges and textures being represented including and replication of the oblique craniotome markings. Visible upon its surface were sub-millimeter ‘layers’ which represented artifacts from the stereolithographic process. On the inner surface, the middle-meningeal groove was replicated in its entirety, and at an appropriate depth of ∼1 mm. Towards the superior arterial bifurcation, the shallow depth of these grooves was only partially replicated. The smaller 3d-scanner derived flap was also an identical replication of the original specimen in terms of shape and size. It measured 48 mm in anteroposterior length (1mm larger than specimen (+2.1%)) and 50 mm in vertical height (1mm smaller than specimen (-2%)). It approximated the real specimen in thickness, measured at its anterior, posterior, superior and inferior edges. Edge thickness was between 2-3 mm. Like the larger part, it possessed build process layer-artifacts but retained an adequate amount of surface detail including a shallow middle meningeal groove on its inner aspect. In comparison, the CT-derived models were notably larger and thicker than the 3D scanned models. The hemicraniectomy was 148mm in anteroposterior length (6.5% larger than the 3D scanner model) and 117 mm in vertical height (9.3% larger than the 3D scanner model). As these parts were manually traced directly from the skull model, they contained a component of the ventrolateral frontal bone (lateral orbital ridge portion). Another notable feature was a reproduction of the bulging lip on the posterior-external surface, i.e. reproduction of the modeling artifact discussed above. This lip, though less evident throughout the rest of the model, continued around the edges, giving them a curved appearance differing from the original. It contributed to a general thickness of 6-7mm at its frontal, superior and posterior aspects. At its most inferior, temporal aspect, it was ∼ 3mm thick. Interestingly, the surface tessellations created by the 1.25 mm slicing of the CT scan were reproduced by the stereolithographic process. The CT-derived smaller flap was 54 mm in length and 61 mm in height, or 8% and 33% larger than the 3D scanner model, respectively. Notably, as the piece was created using a freehand method of virtual extraction, the craniotomy arch at its anteriorinferior corner, from the original specimen and on the scanner derived model, was absent. It had a uniform thickness throughout of ∼ 4mm, furthermore, at its inferior surface and secondary to the freehand method of model creation, was an area of reproduced bony pneumatization. The small piece also contained ridges representative of the 1.25 mm CT slicing and the stereolithographic tessellations giving it a textured appearance. Notably, for both CT derived pieces, artifacts created by metallic screws that had been present in the original specimen for fiducial navigation were reproduced on both inner and outer surfaces. These screws were present for the CT scanning process, but removed for the 3D scanner process, and thus were absent from the latters’ models. The middle meningeal groove was discernible on both pieces, however, it was considerably shallower when compared to the scanner derived pieces. Overall, the CT derived models were larger, thicker, smoother at the edges and more robust than the 3D scanner derived pieces (Figures 4A-D, 5A-B).

**Figure 4 A-D.**
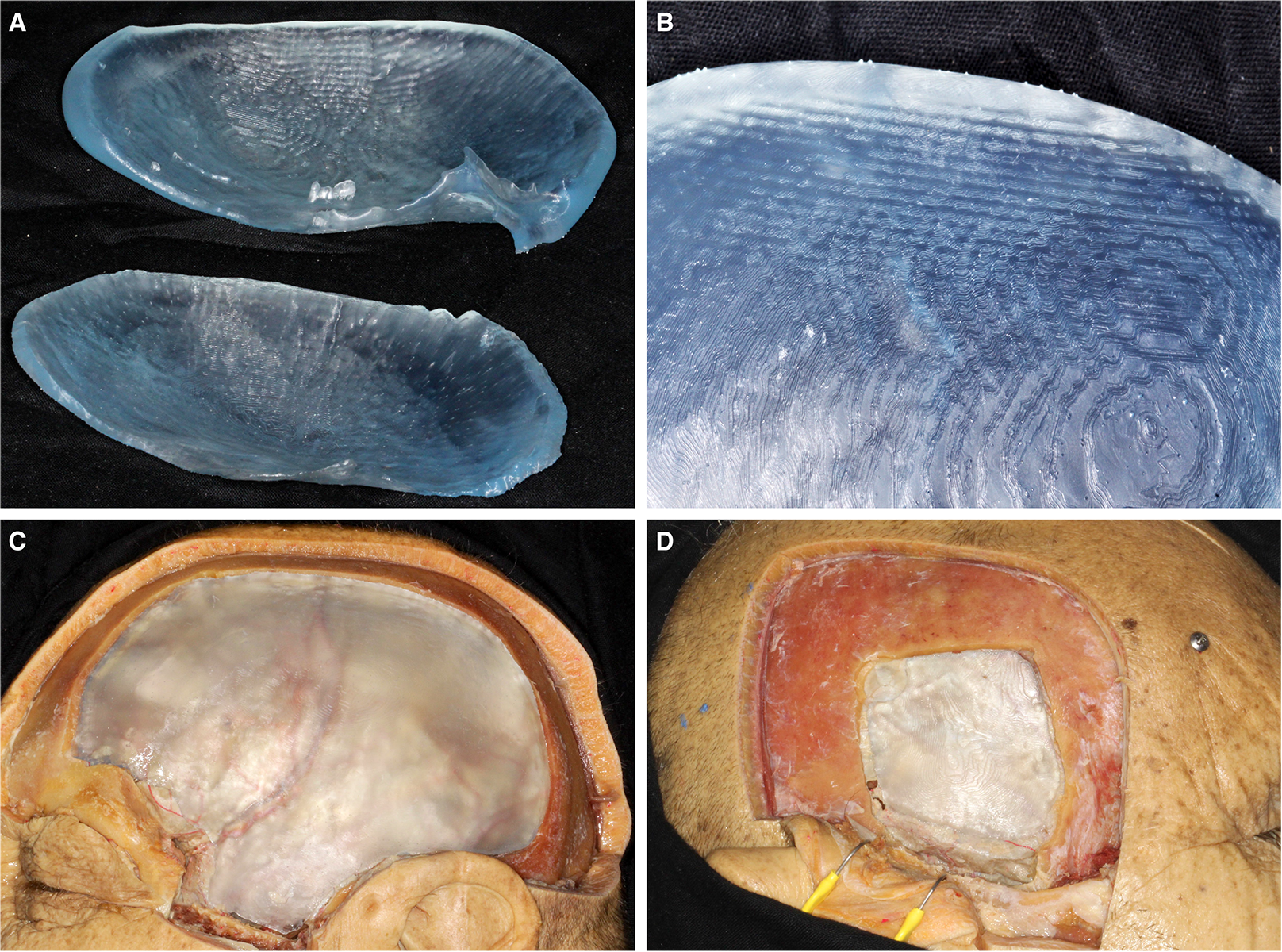
A – External aspects of original hemicraniectomy flap (top middle), 3d-laser scanner model (bottom left) and CT-derived stereolithographically printed flaps. Apparent on the original specimen is the filled defects of the removed screws, the coronal and sphenoid sutures. Note that on the 3D scanner model, the edges and dimensions approximate the original. Note the small defect in the ventral portion of the model, caused by inability for the stereolithographic hardware to build extremely thin models. The CT derived model deviates from the original due to the portion of the ventrolateral frontal bone, and a prominent lip visible on the posterior portion of the flap. B – Internal aspects of the original hemicraniectomy flap (top middle), 3D-laser scanner model (bottom left) and CT-derived stereolithographically printed flaps. Apparent on the original specimen is the filled defects of the removed screws and the middle meningeal groove, along with smaller vascular impressions. Note the edges of the 3D scanner derived piece approximate the edges of the original piece almost identically. This is in comparison to the CT-derived piece, which has curved edges considerably thicker than either the 3D derived model or the original specimen. Also visible on the CT-derived piece are the 2 metallic artefacts from the screws. C - External aspects of original middle-fossa approach flap (top middle), 3d-laser scanner model (bottom left) and CT-derived stereolithographically printed flaps. Apparent on the original specimen is a hole from previous screw placement. Note the consistent size approximation of the 3D model derived piece and the original. This is in comparison to the CT-scanner derived piece, which is substantially larger than either of the other pieces. Also demonstrated on the CT piece is the metallic artefact created from the screw. D - Internal aspects of original middle-fossa approach flap (top middle), 3d-laser scanner model (bottom left) and CT-derived stereolithographically printed flaps. Visible on the original specimen is hole from previous screw placement and the middle meningeal groove, running obliquely. Note the consistent size approximation of the 3D model derived piece and the original. This is in comparison to the CT-scanner derived piece, which is substantially larger than either of the other pieces. Also demonstrated on the CT piece is the metallic artefact created from the screw and the area of bony pneumatisation from the original skull model.

**Figure 5 A-D.**
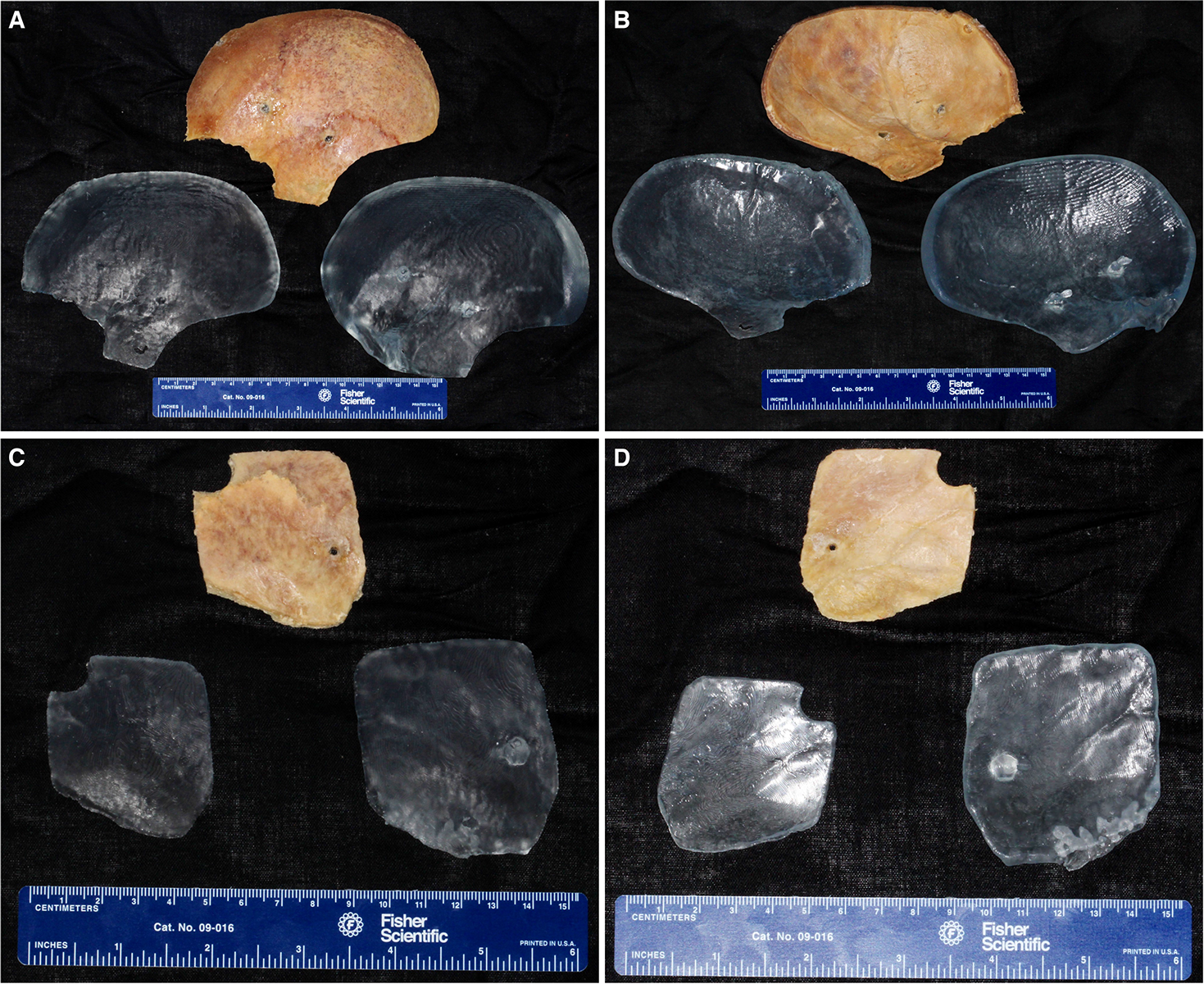
A – An offset view of the CT-derived (top) and laser scanner derived (bottom) stereolithographic pieces. This view is intended to demonstrate the visibility of layers which are artefactual from the 1.25mm fine cut CT scan and the relative difference in edge thickness of the two pieces. B – A close-up view of the CT-derived piece, demonstrating concentric and perpendicular (horizontal) artefacts from the CT scanning process, which were reproduced on the specimen. Visible best on the bottom-left of the image are the ‘tesselations’ which occur due to the sub-millimeter layer-based stereolithographic process. C – A left-lateral view of the cadaveric specimen with implanted 3d-derived stereolithographic hemicraniectomy ‘prosthesis’ placed. Note that the piece fits almost identically to the original. The middle meningeal artery can be discerned to approximate with the groove created on the piece. Also visible are the sub-millimeter tessellations created by the building process. D – A right-lateral view of the view of the cadaveric specimen with implanted 3d-derived stereolithographic middle-fossa approach ‘prosthesis’ placed. Note a ∼1mm gap between the specimen and prosthesis, secondary to material obliteration during craniectomy. Furthermore, the ‘waved’ appearance is representative of the layer-based build process.

Finally, we attempted to fit the models in the craniectomy defects within the specimen. The 3D scanner models fit snugly within the original defect, with ∼ 1 mm of free space between the defect and prosthesis on both the right and left. This gap was due to loss of material obliterated during the craniectomy. The underlying middle meningeal artery lined up with the grooves on the prosthesis on both sides (Figure 5C-D). The CT scanner derived pieces were too large to fit within the defects, primarily due to the lips created around their edges. The pieces could have to be trimmed to fit, which we elected not to do due to the relative rigidity of the build medium.

## 4. Discussion

In this study, we have successfully demonstrated 3 methods of creating virtual 3D models of craniectomy flaps. 2 out of 3 of these methods yielded appropriate prototypes for stereolithographic printing. Our ultimate goal is to use this process to eventually yield clinically viable alloplastic prostheses using simplified methods, and without need for 3^rd^ party outsourcing, provided the necessary knowledge and equipment is available to clinicians.

### 4.1 Virtual Model Creation

Fine-cut CT scanning is a readily available clinical tool, and its intraoperative use in neurosurgery is long established (Butler et al., 1998; Lunsford, Parrish, & Albright, 1984). As it is an axial, X-ray based modality it gives optimal views of bone compared to both standard X-ray and magnetic resonance imaging (MRI). It is therefore an ideal resource to create bony models. In cases of trauma, it is likely that patients will undergo bony-window CT scan during initial assessment. As our models were created using the ‘negative’ i.e. area surrounding the craniectomy, this method lends itself well to cases of compound skull fractures, where the removed material may not yield a single piece of bone. Whilst pre-operative CT scanning is routine and relatively easy to obtain, intraoperative CT scanning may not be, especially with growing use of MRI (Alexander, Moriarty, Kikinis, Black, & Jolesz, 1997). Post-craniectomy CT scanning may therefore entail infection, anesthetic or logistical risks if the patient has to be transferred out of the operating room or neurosurgical unit. As CT scanners are generally a standard piece of medical equipment, equipment acquisition likely entails no extra cost. Once obtained, creation of 3D models is relatively quick and cheap using freely available software, as we have demonstrated. Creating the printable model is a more complex process, however. We utilized superimposition and planar offset of pre-and-post craniectomy scans to create a template outline that could be manually traced to yield the desired shape. There may be other ways to create models from pre- and-post operative scans; however we felt that this was the most intuitive and simplistic method. Other issues that may arise during model creation, and as we experienced, were the presence of tomographic artifacts, i.e. the metallic screws. Metallic shrapnel may therefore complicate the scanning process during cases of penetrating trauma, injury or device implants. We found the artifacts impractical to virtually remove without damaging the surface of the pieces and elected to leave them in. Despite these issues, virtual model creation was a relatively quick and free process.

The models created using the 3D light scanner were ultimately unfeasible for stereolithographic printing due to numerous surface defects and alignment issues. These defects likely arose due to the relatively high number of individual scans that comprised the final virtual objects. Despite the supplied DAVID 3D software having the ability to automatically register and align the individual scans, we found that this alignment was suboptimal, and required considerable manual manipulation of single scans to yield what appeared to be an acceptable model. A potential reason for the lack of success of this method was that the bone pieces were laid flat upon the turntable during scanning. These pieces had negligible vertical height and complex edges. Furthermore, we postulate, that due to laying them flat on the turntable, the false surface texture may have been secondary to shadows cast by the projector during acquisition. Scanning time is a further issue. Each model required several hours to acquire, align and create. Overall this method proved to be the most time consuming out of the 3. Due to low cost and high mobility, the DAVID 3D camera and turntable may be practical for rapid transport and clinical use, but further work is necessitated to determine its potential applications.

The FARO Laser ScanArm provided the best quality virtual and real models out of the 3 methods. It represented the most rapid and least convoluted solution for scanning the removed bone pieces. We are not aware of any previous applications of this technology to 3D printing in neurosurgery. The design of this system is ideal for surgical application, as it bears range and axial movement similar to surgical microscopes. Our study shows that it could be used intraoperatively to either scan removed hard or soft tissue, or the defect that has been created. For infection control, a simple plastic cover may be placed over it, as is the case for surgical microscopes or X-ray equipment. Furthermore, this arm-based laser scanner can potentially be mounted on a trolley or fixed within an operating suite. The main drawback of using the laser 3D scanner is its cost, which is substantially higher than the DAVID 3D light-scanner and CT scanning. Secondly, the software required to run the 3D laser scanner is a dedicated CAD solution, specific to engineering applications and which would entail substantial cost and a learning curve for clinicians. As such, further clinically beneficial applications of 3D scanning with this device in neurosurgery should be identified to justify the cost of acquiring relevant hardware.

#### 4.2 3-Dimensional Printing

Regarding 3-dimensional or stereolithographic printing technology, relevant neurosurgical applications include 1.) the creation of implantable synthetic prostheses 2.) creation of patient-specific medical models for use in pre-operative planning or radiotherapy (e.g. headframes) and 3.) the creation of 3D teaching aids for training of students, residents or fellows.

Perhaps the most exciting potential application of stereolithographic printing in medicine is for creating patient-specific synthetic prostheses (Klein et al., 2013; Randazzo et al., 2016; Tomasello et al., 2016). This study aimed to create a simplified workflow for constructing implantable cranioplasty flaps; from data acquisition, virtual modeling, and eventual physical production. Moreover, we aimed to create a process that could be conducted entirely by the clinician, with minimal third-party input. Alloplastic skull prostheses should serve both structural (discussed in 4.3) and cosmetic functions. Pertinent to cosmetics is the closure of craniectomy defects and reconstruction of the traumatically damaged skull or maxillofacial bone. The appeal of CAD in cosmetic reconstruction is obvious, as it allows the clinician maximal scope of tailoring prostheses specifically to the individual. This approach can save the time required and number of procedures, ultimately resulting in superior cosmetic outcomes (Chae et al., 2015; Gerstle, Ibrahim, Kim, Lee, & Lin, 2014). As such, the application of 3D aided CAD processes in craniofacial procedures has been explored in the literature, and virtual modeling is actively utilized in the design and creation of implantable prostheses (Orentlicher, Goldsmith, & Horowitz, 2010; Pham, Rafii, Metzger, Jamali, & Strong, 2007). Limitations at present appear to be cost and access to both hardware and software required to design and build the prostheses, and subsequent training amongst clinicians. Furthermore, the issue of integrating 3D CAD technology with FDA approved synthetic materials provides a significant barrier to widespread institutional implementation (Chia & Wu, 2015; Murphy, Skardal, & Atala, 2013).

The use of stereolithographic printing in medical modeling is already an accepted practice (McGurk et al., 1997; McMenamin, Quayle, McHenry, & Adams, 2014; Rengier et al., 2010). It allows creating 3D models of anatomical structures unique to the subject and their pathology. These models can enhance clinician understanding of complex anatomical structures prior to surgery by providing them with spatial information superior to that of radiographic data alone. Furthermore, the stereolithographic equipment and printing would not require the rigorous approval processes required for implants. It can thus be conducted relatively cost-effectively. A potential drawback, however, is the need to justify this cost of equipment acquisition, materials and time required for model creation. As the models would be patient-specific rather than generalized, they would be single-use only. Moreover, this technique may be limited to modeling of relatively small anatomical structures due to logistical and cost considerations of printing larger objects. A potential area for exploration is 3D printing in skull-base surgical planning, as this is an area possessing complex spatial anatomy (Stadie et al., 2008; Waran, Narayanan, Karuppiah, Owen, & Aziz, 2014).

### 4.3 Materials and Structural Considerations

As previously mentioned, the greatest limitation to widespread adoption of institutional, clinician-operated 3D printing for alloplastic implant creation is lack of integration of the technology and a selection of safe, FDA-approved structural materials. These materials should appropriate or exceed structural properties relative to the native material, e.g. bone whilst causing minimal immunoreactivity and an effective barrier to pathogens. These materials may also promote tissue regeneration whilst providing a structural scaffold (Cox, Thornby, Gibbons, Williams, & Mallick, 2015; Leukers et al., 2005; Park, Lee, & Kim, 2011). Specific to cranioplasties are the physical strength of the material and its ability to enhance cosmesis (Aydin et al., 2011). There is no ‘gold standard’ in choice of synthetic cranioplasty material, with synthetic polymers including methacrylate (Cooper, Schechter, Jacobs, Rubin, & Wille, 1977; Findler, Sela, & Sahar, 1979), PEEK (Ng & Nawaz, 2014; O’Reilly et al., 2015), in addition to metals (Hill, Luoma, Wilson, & Kitchen, 2012; Stoodley, Abbott, & Simpson, 1996) and ceramics (Kobayashi, Hara, Okudera, Takemae, & Sugita, 1987; Miyake, Ohta, & Tanaka, 2000) being used. Each has its own advantages and disadvantages (Cabraja, Klein, & Lehmann, 2009; Jaberi, Gambrell, Tiwana, Madden, & Finn, 2013; Moreira-Gonzalez, Jackson, Miyawaki, Barakat, & DiNick, 2003; Rosenthal et al., 2014). As the methacrylate medium is commonly mixed and applied during surgery, 3D printing has been used to create molds (Chiarini et al., 2004; Lee, Wu, Lee, & Chen, 2009). This, may be a less intuitive approach considering available technology to directly print implants. In some instances, a titanium mesh is placed as a structural foundation prior to methacrylate application (Blum, Schneider, & Rosenthal, 1997). Direct metal laser sintering is a method to create complex 3D objects from powdered alloys of titanium (Ciocca, Fantini, De Crescenzio, Corinaldesi, & Scotti, 2011; Mazzoli, 2013). The laser locally heats a metallic powder to melting point, ultimately causing fusion of the materials in a successive layer based process. In addition to solid planar objects, meshes and even chain-linked structures can be manufactured by this technique. Potential application of this technology in cranioplasties is to create a metallic foundational base prior to methacrylate application. Alternatively, metallic cranioplasties may be created in their entirety using this process, and has been demonstrated (Jardini et al., 2014). As titanium is already widely used for prosthetics, little to no prior clinical approval would be required for creation of these implants provided they were appropriately finished, sterilized and affixed. PEEK is structurally similar to native bone and is already widely used in neurosurgical and craniofacial procedures. In addition, it is lightweight, radiolucent and can be sterilized repeatedly (Aydin et al., 2011; Ng & Nawaz, 2014; O’Reilly et al., 2015). Currently, clinical use of this technology entails recruitment of off-site third parties, potentially creating confidentiality, geographical and time constraint issues for the clinician.

The strength of synthetic materials is particularly relevant in orthopedic or spinal procedures (Lethaus et al., 2012), whereby prosthetic implants likely a load bearing function. It is therefore particularly important for the physical properties of the construction materials to be equal or superior to those of bone. Naturally, metallic mediums, such as titanium, lend themselves to the creation of structural prostheses. We are aware of at least one clinically reported, specially manufactured, 3D printed titanium prosthesis used for cervical vertebral replacement in a pediatric patient, with no major complications at 1-year followup (Xu et al., 2016). Due to the complexity of creating structurally viable, 3D printed synthetic materials, considerable further research is required.

## 5. Conclusions

In this study, we have successfully explored the concept of clinician-created virtual and physical models of cranioplasty prostheses. Though we did not create the models in a clinically viable, implantable synthetic medium, the models served to demonstrate the methodology, which may ultimately be utilized to create neurosurgical prostheses in an FDA-approved synthetic plastic or metallic medium. Furthermore, we emphasize a user-based approach, allowing creation of alloplastic implants on-site and in a rapid timeframe. Currently the greatest barriers to widespread adoption of this technology are the cost of equipment, lack of knowledge and training amongst clinicians and introduction of commercial, FDA-approved mediums for printing. We conclude by calling for greater research into this method.

## 6. Acknowledgements

We would like to thank J. Andrew Holmes at the ANSYS Additive Materials and Manufacturing Laboratory at the Swanson School of Engineering, University of Pittsburgh and Wendy Fellows-Mayle PhD, at the Surgical Neuroanatomy Lab, University of Pittsburgh for their assistance with this project.

## 8. Ethics

This study was approved by the Institutional Review Board at the University of Pittsburgh.

